# Native in situ architecture of the human inactive X chromosome revealed by correlative light and electron microscopy

**DOI:** 10.64898/2026.09.04.749547

**Authors:** Barbara Hübner, Mei Ling Wee, Harvey Yin Seng Chan, Sara Sandin, Alexander Ludwig

## Abstract

Knowledge of the nuclear organization of specific chromatin domains at the ultrastructural level remains limited. To address this, we employed correlative light and electron microscopy (CLEM) approaches to investigate the inactive X chromosome (Xi) in female human RPE1 cells. Tandem fusion of macroH2A (mH2A), a histone variant enriched on the Xi, to eGFP and APEX2 enabled direct labeling and identification of the Xi by light microscopy, transmission electron microscopy, and electron tomography using conventional CLEM. The eGFP tag further allowed us to visualize the Xi at single nucleosome resolution under close-to-native conditions using cryo-CLEM. We found that, although heterochromatin domains of the Xi are substantially larger than those of autosomal regions, their chromatin density was not significantly different. Furthermore, our data reveal the organization of the interchromatin compartment (IC) within the Xi, characterized by a network of larger lacunae and smaller channels. Overall, we provide an ultrastructural characterization of a specific heterochromatin domain at unprecedented resolution, advancing our understanding of 3D chromatin organization and nuclear architecture *in situ*.

**SIGNIFICANCE:** How chromatin is organized within the nucleus is a fundamental question in cell and molecular biology. Recent cryo-electron tomography studies have revealed chromatin and nucleosome organization in situ but lack molecular identification of the chromatin domains examined. Here, we investigate the ultrastructure of the human inactive X chromosome (Xi), or Barr body, a highly compact heterochromatin domain, in female RPE1 cells. By combining conventional and cryogenic correlative light and electron microscopy (cryo-CLEM), we characterize its three-dimensional architecture under near-native conditions and resolve chromatin to the level of individual nucleosomes. Our findings reveal the ultrastructure of a defined heterochromatin domain in situ and establish a framework for connecting chromatin molecular identity with native three-dimensional architecture.

## INTRODUCTION

Chromatin structure is a key regulator of gene silencing in eukaryotes (Minami et al., 2026; Paldi and Cavalli, 2026). Despite decades of research, how nucleosome arrays are organized into higher-order chromatin domains within the nucleus remains poorly understood and highly debated (Liu et al., 2024; Maeshima, 2025; Hou and Zhang, 2025). The inactive X chromosome (Xi) provides a particularly striking model for studying highly compacted chromatin. The Xi results from the random inactivation of one of the two X chromosomes in female mammalian cells to ensure dosage compensation of X-linked gene expression between both sexes (Lyon, 1961; Galupa and Heard, 2018; Loda et al., 2022). The resulting Xi, and particularly its more compacted core historically termed the “Barr body” (Barr and Bertram, 1949), exhibits a distinct chromatin architecture compared with the active X chromosome (Xa) and autosomes (Galupa and Heard, 2018; Loda et al., 2022). X chromosome inactivation (XCI) is initiated and maintained through a complex interplay of factors, with the long noncoding RNA XIST (X-inactive specific transcript) playing a central role in establishing transcriptional silencing across much of the chromosome (Jacobson et al., 2022; Penny et al., 1996; Galupa and Heard, 2018; Loda et al., 2022; Martitz and Schulz, 2024). This process is accompanied by extensive reorganization of the Xi at multiple levels, including its chromatin domains (Nora et al., 2012; Rao et al., 2014; Deng et al., 2015; Minajigi et al., 2015; Darrow et al., 2016; Giorgetti et al., 2016), chromatin interactions (Splinter et al., 2011; Nora et al., 2012; Rao et al., 2014; Deng et al., 2015; Darrow et al., 2016) and chromatin compartments (Rao et al., 2014; Darrow et al., 2016; Miura et al., 2018; Bauer et al., 2021; Martitz and Schulz, 2024). These features have been extensively characterized using molecular and chromosome conformation capture (3C)-based approaches, providing important insights into the genomic organization of the Xi. However, how these molecularly defined domains and compartments are physically arranged at the nanometer-to-micrometer scale within the nucleus remains much less well understood.

Super-resolution fluorescence microscopy has provided important insights into nuclear organization (Birk, 2019; Burgers and Vlijm, 2023; Cremer et al., 2015). Chromosome territories are permeated by the interchromatin compartment (IC) which is largely free of chromatin and carries splicing speckles and nuclear bodies. Along with the perichromatin region (PR) that contains decondensed euchromatin and harbors macromolecules involved in many nuclear functions, the IC forms the active nuclear compartment (ANC) (Cremer et al., 2020). Using three-dimensional structured illumination microscopy (3D-SIM), Smeets and coworkers (Smeets et al., 2014) confirmed previous observations from conventional electron microscopy (EM) (Rego et al., 2008) that the ANC was significantly reduced but not absent in the Xi. Consistent with its heterochromatic nature, chromatin domains in the Xi were suggested to be more closely packed than those in autosomal regions (Galupa and Heard, 2018; Loda et al., 2022; Martitz and Schulz, 2024). However, the ∼100–130 nm lateral and ∼250–340 nm axial resolution of 3D-SIM is insufficient to resolve the ultrastructural organization of chromatin and the IC (Schermelleh et al., 2010). Conventional EM studies of the Xi have so far provided mainly low- to intermediate-magnification projection images (Rego et al., 2008; Bourgeois et al., 1985; Baghestani, 2015), leaving its three-dimensional ultrastructure largely unexplored. Moreover, both fluorescence microscopy and conventional EM typically require chemical fixation and other sample-processing steps that can alter cellular ultrastructure through cross-linking, material loss, or shrinkage (Mollenhauer, 1993; McDonald and Auer, 2006). Cryo-EM methods overcome many of these limitations by preserving cellular structures under close-to-native conditions, providing a powerful approach for resolving cellular ultrastructure at high resolution (Wagner et al., 2017; Zheng and Cai, 2024; Hou and Zhang, 2025; Kechagia and Medalia, 2026; Chen et al., 2026).

Here we established two complementary workflows to image the Xi in female human retinal pigment epithelial (RPE1) cells using correlative light and electron microscopy (CLEM). To selectively label the Xi, we fused macroH2A (mH2A), a histone variant enriched in this chromosome (Costanzi and Pehrson, 1998; Chadwick and Willard, 2001; Costanzi and Pehrson, 2001), to a tandem eGFP-APEX2 tag. This allowed us to identify the exact sub-nuclear localization of the Xi by light microscopy (LM) and subsequently study its ultrastructure by transmission electron microscopy (TEM), electron tomography (ET) and cryo-ET. APEX2 with conventional EM sample preparation was used to specifically enhance the contrast of chromatin within the Xi to achieve direct labeling for EM. In addition, cryo-ET of vitrified, unstained cells served to resolve the native architecture of the Xi down to the single nucleosome level. Our tomography data provides ultrastructural details of Xi-linked heterochromatin *in situ* at unprecedented resolution.

## RESULTS and DISCUSSION

### Conventional CLEM with APEX2 allows direct labeling of the Xi for LM and EM

To specifically label the Xi for CLEM, we fused mH2A in tandem to eGFP and APEX2. APEX2 is a modified peroxidase, which can provide a selective and electron-dense label for EM (Martell et al., 2012; Lam et al., 2015) and is suitable to identify specific chromatin domains (Hübner et al., 2022). This approach allowed us to directly localize and sequentially visualize the Xi in the same cells in LM via the eGFP signals as well as in TEM and ET via the APEX2 probe (Fig 1A-B). After the APEX reaction and osmium staining, dark contrast was visible in the EM images corresponding to the sites of the eGFP signals. In many cases the APEX signals clearly touched the nuclear envelope (NE), contradicting previous observations that the Xi is separated from the NE by a layer of autosomal chromatin (Baghestani, 2015) (for simplicity, we include the Xa into the terms ‘autosomal’ and ‘autosomes’, since its structure, organization and appearance is similar to autosomes (Galupa and Heard, 2018; Smeets et al., 2014)). Multiple larger and smaller areas of low contrast were visible within the Xi, similar to the areas of low DNA density observed with fluorescence super-resolution microscopy (Smeets et al., 2014).

**Figure 1:**
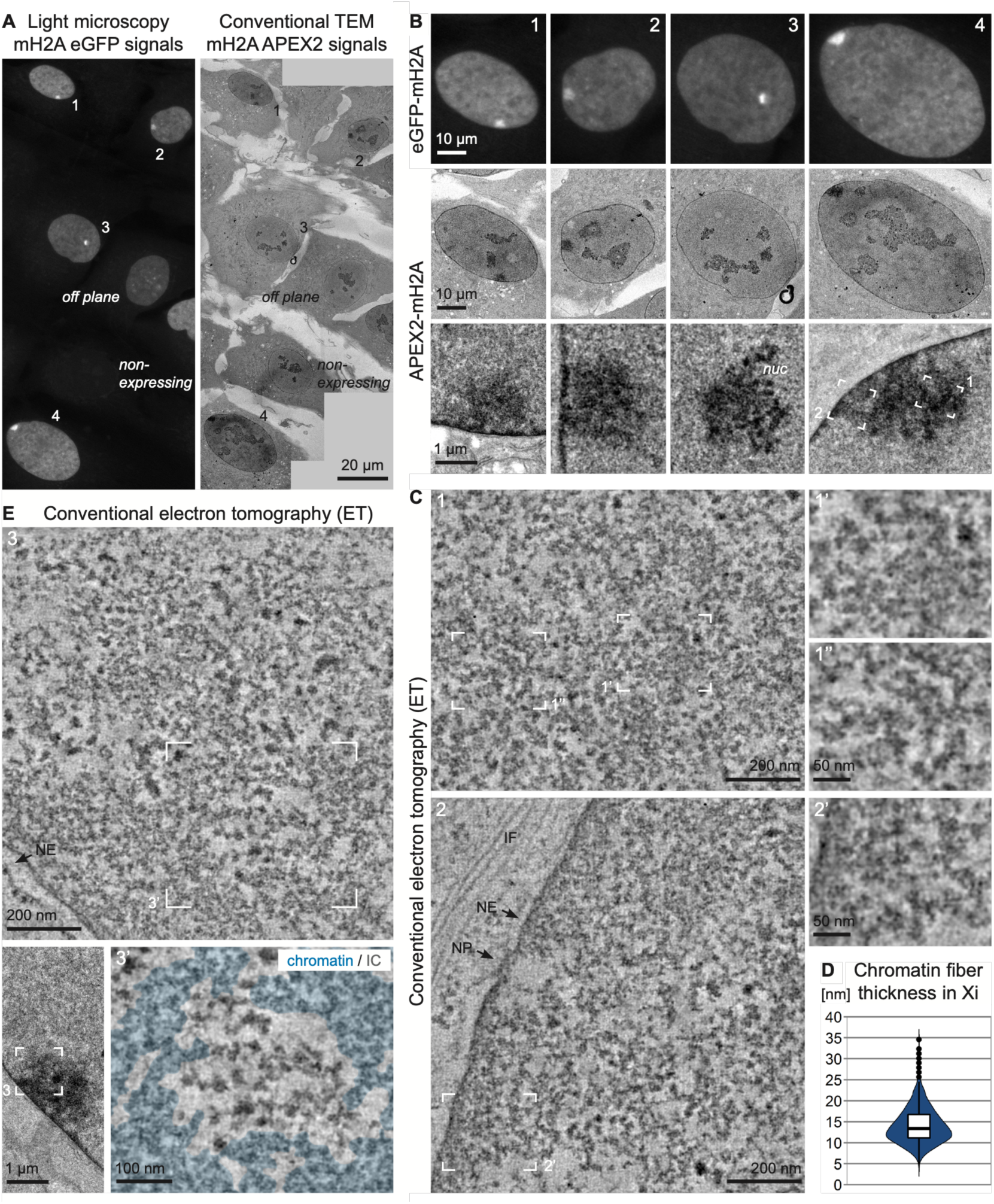
Conventional CLEM and ET using APEX2 allows direct identification of the Xi in LM and EM. **(A)** Confocal light optical section of eGFP signals of RPE1 cells stably expressing eGFP-APEX2-mH2A (left) and corresponding tile montage of TEM images after the APEX2 reaction and processing for EM (right). A non-expressing cell and a cell where the Xi is off plane is included. **(B)** Magnification of cells 1-4 from (A). The Xi is further magnified in the bottom row. nuc = nucleolus. **(C)** ET average slices (combined thickness of 5.6 nm) of the boxed areas of cell 4 in (B). Subareas showing heterochromatin are further magnified on the right. NP = nuclear pore. NE = nuclear envelope. We regularly (33%) found bundles of intermediary filaments (IF) in the cytoplasm adjacent to the Xi (example 2) and occasionally (16%) also at autosomal regions (SFig 2C, right) (n = 24 / 19 for the Xi / autosomal regions, respectively, APEX and cryo data combined). **(D)** Evaluation of chromatin fiber thickness in the Xi. Violin plot. The median (13.4 nm) and the interquartile range are indicated in the overlaid box plot. n = 2110 measurements from 11 tomograms of 10 cells. **(E)** Top: ET average slice (5.6 nm) of the boxed area of another cell shown on the bottom left. A subarea highlighting an IC lacuna is further magnified on the bottom right (chromatin is overlaid with cyan).

To validate our imaging workflow, we analyzed mH2A-eGFP-APEX2 expressing cells using confocal microscopy (SFig 1). First, we confirmed in mitotic cells that indeed only one chromosome (i.e. the Xi) showed the enrichment of mH2A signals (SFig 1A). We then investigated whether the mH2A-labeled Xi contained a dense DNA core (Barr body) (SFig 1B). Consistent with previous observations (Bonora and Disteche, 2017), not all cells showed a clear Barr body (BB) upon DNA staining with Hoechst: in 26% of cells the BB was barely visible and 7% of cells did not show any apparent chromatin compaction at the site of mH2A accumulation (SFig 1B, top). In addition, a small percentage of cells (5%) did not show an enrichment of mH2A at a specific site in the nucleus, despite expressing the mH2A construct (SFig 1B, bottom), in line with previous reports (Costanzi and Pehrson, 1998; Chadwick and Willard, 2002). Regarding the structure of the Xi, we found that 21% of cells (SFig 1C) exhibited a bipartite appearance of the mH2A signals (asterisks in SFig 1B), matching earlier light microscopy data (Crouch and Barr, 1954; Klinger, 1958) and 3C-based methods demonstrating that the Xi is organized into two mega- or superdomains (Rao et al., 2014; Deng et al., 2015; Minajigi et al., 2015; Darrow et al., 2016; Giorgetti et al., 2016). Further, mH2A signals always (100%) extended over the BB as revealed by Hoechst staining, in the majority of cases (78%) even clearly (SFig 1D-E). This is consistent with previous observations that the BB is smaller than the Xi chromosome territory (Clemson et al., 1996; Hall and Lawrence, 2010; Teller et al., 2011). We found the Xi to be located at the nuclear envelope (NE) in 58% of cells when evaluated in 2D and in all cells (100%) when evaluated from 3D image stacks (SFig 1F). Further, this chromosome was associated with a nucleolus in around 40-65% of cells (SFig 1G-I). These results match previous in-depth analyses (Bourgeois et al., 1985; Rego et al., 2008) and numerous additional studies going back to the early days of Xi-related research.

Overall, our observations are in agreement with previously published results, indicating that expression of eGFP-APEX2-tagged mH2A does not interfere with Xi behavior or basic structure.

### Electron tomography (ET) provides three-dimensional insight into the architecture of the Xi

Next, we used ET to study the chromatin organization and architecture of the Xi at high resolution (Fig 1C-E, SFig 2). As already indicated from the projection TEM images (Fig 1B bottom, SFig 2A, D), ET confirmed that the edges of the APEX-labeled chromatin domain were irregular and contrast usually faded into the surrounding autosomal chromatin (Fig 1E, top part of the tomogram, SFig 2 example 1, 4, 5). This suggests that the Xi is not spatially isolated but embedded in and in contact with other chromatin, despite its particular, largely heterochromatic and transcriptionally inactive state. Chromatin inside the Xi (Fig 1C) largely appeared as small clusters or irregular fibers with variable conformations. Fiber width ranged from 4.5 nm up to 34.6 nm with a median of 13.4 nm (Fig 1D). These measurements are comparable to what we previously found in telomeric chromatin of MEFs (Hübner et al., 2022). Occasionally we observed small patches where the interchromatin space was reduced, both internal in the nucleus / Xi (e.g. Fig 1C inset 1’, right half) as well as at the NE (e.g. Fig 1C inset 2’, left half). Whether these areas are built up from several fibers that cannot be resolved or rather represent one broad “fiber” or domain with a more intermingled chromatin organization cannot be answered from our data.

**Figure 2:**
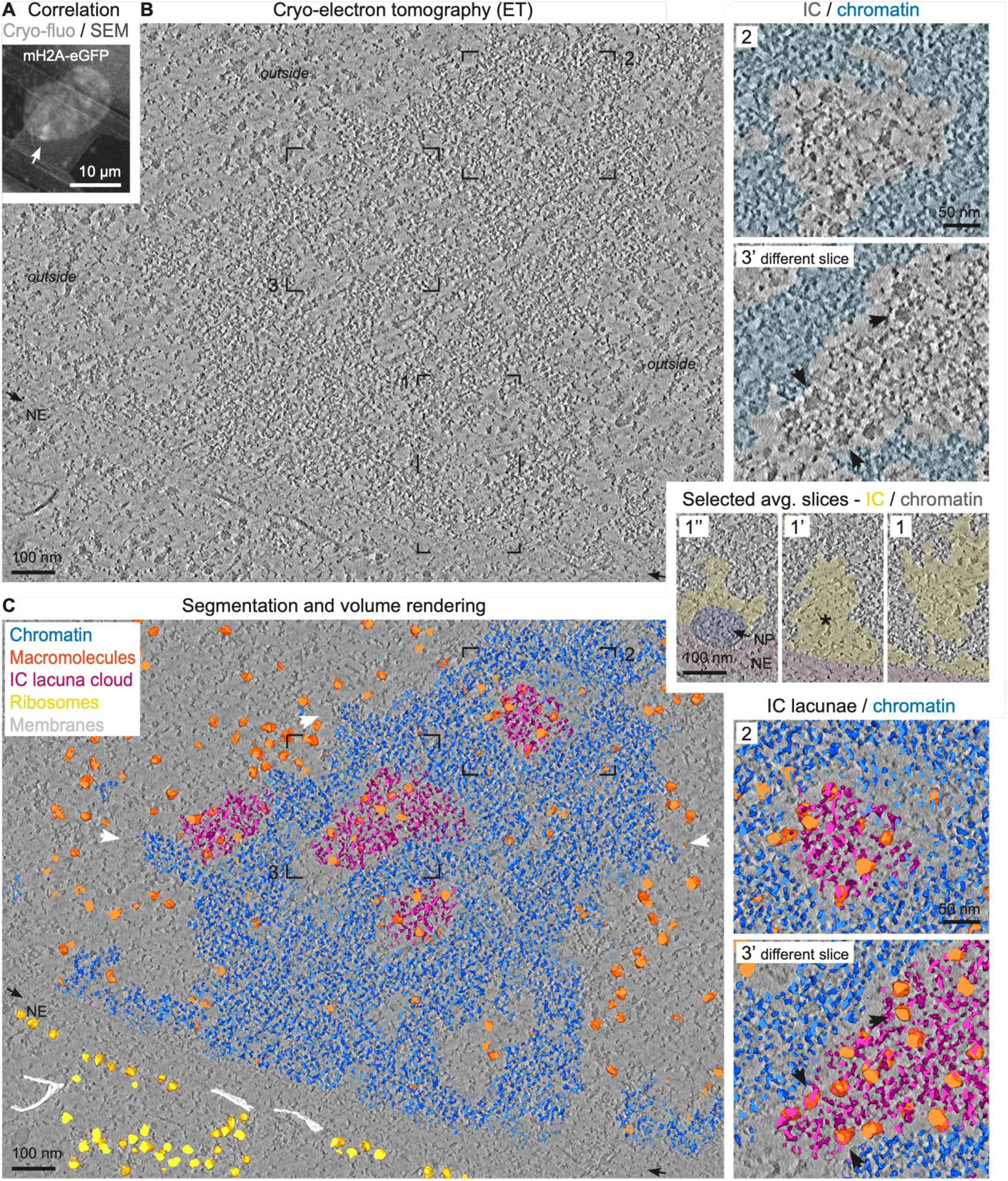
Cryo-CLEM and ET of the Xi under close to native conditions and segmentation and volume rendering. **(A)** The Xi was located on the lamella based on the correlation of cryo-fluorescence signals of eGFP-mH2A (confocal spinning disk mode, maximum intensity projection) with SEM images after completed milling (top: cryo-fluorescence alone, bottom: overlay). See also SFig 3A-B. **(B)** Cryo-ET average slice (combined thickness of 5.2 nm; VPP, K2, -0.5 µm defocus, 33k×) of the Xi of the cell shown in (A). Nucleoplasm outside the heterochromatin domain of the Xi comprising the IC and euchromatin (EC) is marked with ‘outside’. The black boxed areas are further magnified on the right. In inset 1 and the two additional average slices (1’ and 1’’) selected from the volume the IC/EC is overlaid with yellow. The asterisk highlights an IC lacuna below the nuclear pore. In inset 2 and 3’ (which comes from a different average slice but the same location as box 3) chromatin is overlaid with cyan. Arrowheads in 3’ point at macromolecular complexes of IC lacunae inside the Xi. NE = nuclear envelope (orientation indicated by black arrows in (B), overlaid with magenta in inset 1). NP = nuclear pore, overlaid with purple. **(C)** Segmentation and volume rendering corresponding to Fig 2B comprising a thickness of 20.7 nm (the corresponding data comprising a thickness of 112.3 nm is shown in SFig 4B). A single tomogram slice is shown in the background (gray). The boxed areas are further magnified on the right (inset 3’ comes from a different average slice but the same location as box 3; the magnification of box 3 is included in SFig 4B). The white arrowheads point at heterochromatin extensions of the Xi connecting to euchromatic regions. NE = nuclear envelope (orientation indicated by black arrows). Blue = chromatin, orange = macromolecules, magenta = IC lacuna cloud, yellow = ribosomes, white = membranes. Note that the NE could not be segmented due to lack of contrast as a consequence of being cut at an angle.

Furthermore, we observed expanded regions with increased interchromatin space that contained structures distinct from chromatin that were mostly globular in shape and seemingly more electron dense (Fig 1E inset 3’). These areas usually matched with the areas of low contrast in the TEM images (especially clear in Fig 1B-C example 2 and SFig 2A-B example 2) and represent interchromatin compartment (IC) lacunae. Lacunae – as well as smaller IC channels – are not always easy to identify and often the distinction between chromatin and other macromolecular complexes is open to interpretation. Nonetheless, our data clearly illustrate – for the first time at nanometer resolution – that the IC starts at nuclear pores (NP) (Fig 1C, SFig 2E) and pervades the entire Xi with a network of larger lacunae and smaller channels. For comparison, we also imaged nuclear regions outside the Xi (autosomal regions) (SFig 2C). As expected, and matching with a mainly euchromatic / open chromatin organization of the majority of autosomal regions, most chromatin was much more dispersed in these tomograms and even more difficult to distinguish from the considerably more extensive IC. Also here, we regularly observed dense macromolecular complexes similar to those in the IC lacunae within the Xi.

Taken together, APEX2-tagged mH2A enabled selective contrast enhancement of the Xi, allowing its precise identification in cell sections and high-resolution imaging of its architecture by ET. This approach provides direct three-dimensional ultrastructural insight into the chromatin organization of the inactive X chromosome.

### Cryo-CLEM resolves the native organization of the Xi and its extensive interchromatin compartment (IC)

To image the Xi under close-to-native conditions, we employed cryo-CLEM (Mahamid et al., 2016; Wagner et al., 2017; Zheng and Cai, 2024). In cryo-EM, cells are immobilized by vitrification (rapid freezing to avoid the formation of ice crystals), which maintains the sample in a fully hydrated state and thereby avoids artifacts of conventional TEM sample preparation methods. Our cryo-CLEM workflow involves cryo-fluorescence imaging to identify the Xi in vitrified cells based on eGFP signals, followed by FIB-SEM (focused ion beam - scanning EM) milling (Wagner et al., 2017) to create thin lamellae suitable for cryo-ET (see Methods and SFig 3A-B). We used fiducial marker beads (Arnold et al., 2016; Klumpe et al., 2021; Bieber et al., 2021) and a two-step cryo-LM and -EM correlation approach to determine the precise location of the Xi for EM: Correlation was performed before thinning in order to define the final lamella position for FIB-SEM (SFig 3A, left and middle) as well as afterwards to pinpoint the location of the Xi on the lamella for cryo-ET data collection (SFig 3A right, SFig 3B and Fig 2A). Based on this post-milling correlation 79% of our lamellae contained the Xi.

**Figure 3:**
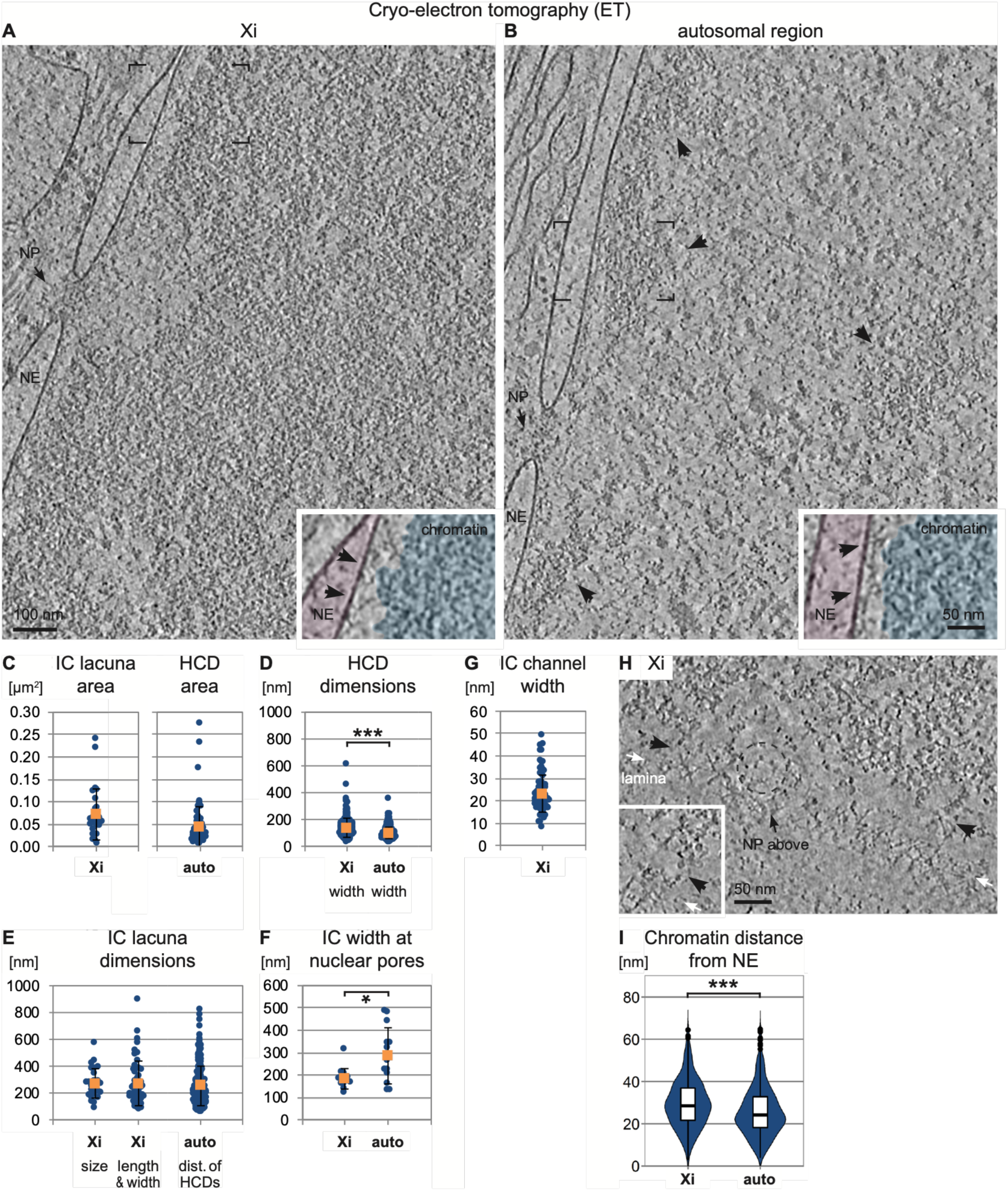
Nuclear architecture within the Xi is distinct from autosomal regions. (A, **B)** Cryo-ET average slices (combined thickness of 5.2 nm; K2, -6 µm defocus, 33k×) of the Xi (A) and an autosomal region (B). NE = nuclear envelope. NP = nuclear pore. Arrowheads in (B) point at heterochromatin domains. The boxed areas are magnified in the insets after angular correction of the tomogram. Chromatin overlaid with cyan. NE = nuclear envelope, overlaid with magenta. Black arrowheads point at contacts between chromatin and the lamina. **(C-G)** Evaluations. Blue = individual values. Orange square = mean. Error bars represent standard deviation. (C) IC lacuna area and HCD area within the Xi and autosomal regions (auto), respectively. Mean = 0.072 µm^2^ / 0.042 µm^2^. n = 26 / 74 measurements from 10 / 18 tomograms of 10 / 13 cells. (D) HCD dimensions (width) within the Xi and in autosomal regions (auto). Mean = 138 nm / 101 nm. n = 247 / 229 measurements from 10 / 18 tomograms of 10 / 12 cells. Statistical significance based on a two-sided *t*-test: p < 0.001. (E) IC lacuna dimensions within the Xi (size = average of length and width) and distance of heterochromatin domains (HCDs) in autosomal regions (auto). Mean = 271 nm / 271 nm / 257 nm. n = 26 / 52 / 216 measurements from 10 / 10 / 20 tomograms of 10 / 10 / 13 cells. (F) IC width at nuclear pores within the Xi and in autosomal regions (auto). Mean = 185 nm / 288 nm. n = 13 / 13 measurements from 9 / 9 tomograms of 9 / 7 cells. Statistical significance based on a two-sided *t*-test: p = 0.014. (G) IC channel width within the Xi. Mean = 23.0 nm. n = 87 measurements from 3 tomograms of 3 cells. **(H)** Cryo-ET average slices (combined thickness of 5.3 nm; F4i, -4 µm defocus, 53k×) of the nuclear periphery of the Xi. The orientation of the lamina is indicated by white arrows. **(I)** Evaluation of chromatin distance from the NE within the Xi and in autosomal regions (auto). Violin plots. The median values (28.5 nm / 24.2 nm) and the interquartile ranges are indicated in the overlaid box plots. Outliers > 65 nm were excluded from the analysis. n = 365 / 441 measurements from 6 / 10 tomograms of 6 / 8 cells. Statistical significance based on a two-sided *t*-test: p < 0.001.

Fig 2B shows cryo-ET data of the Xi collected with the Volta phase plate (VPP) (Danev et al., 2014; Fukuda et al., 2015) along with its segmentation and volume rendering in Fig 2C, SFig 4B and MovieS1. Compared to the APEX data (Fig 1E, SFig 2), the borders of the heterochromatin domain of the Xi appeared more distinct in cryo and stood out more clearly from the adjacent nucleoplasm (labeled with “outside”) (Fig 2B). Nonetheless, the surface of the Xi was still irregular with frequent protrusions, suggestive of contact points with other chromatin regions (Fig 2C, white arrowheads). The IC was also easier to distinguish in cryo compared to conventional TEM. Again, we saw that the IC network started and expanded underneath the NPs and pervaded the entire Xi with larger lacunae and smaller channels (Fig 2B inset 1, SFig 4A, IC overlaid with yellow), which is in agreement with previously published studies (Rego et al., 2008; Smeets et al., 2014; Cremer et al., 2020). In the tomograms, we see multiple small channels that form a continuous network of the larger IC lacunae within the Xi as well as with the surrounding nucleoplasm (SFig 4A, arrowheads). While some lacunae, especially those at the NPs (e.g. Fig 2B inset 1’, SFig 4A, asterisks), contained only few, rather small macromolecules, most harbored large, spherical or slightly elongated assemblies. These were embedded in a mesh-like cloud, reminiscent of chromatin but with a distinct ultrastructure separated from the surrounding chromatin (overlaid with blue) by a rim of extended interchromatin space (Fig 2B-C inset 2, 3’). Interestingly, these assemblies, ∼21 nm in diameter (SFig 4C), were often arranged in rows or chains (Fig 2B-C inset 3’, arrowheads). As we observed similar macromolecules also outside the Xi (Fig 2C, orange), we speculate they represent components of functional nuclear processes such as splicing, replication or transcription. The elongating form of RNA polymerase II can be present in the IC lacunae of the Xi in RPE1 cells (Smeets et al., 2014) and genes escaping X chromosome inactivation were located also within the Xi territory (Teller et al., 2011; Calabrese et al., 2012). Alternatively – because the macromolecules outside the Xi were rarely accompanied by a mesh – the assemblies in the Xi lacunae might represent a specialized feature of the Xi such as Xist foci (Markaki et al., 2021; Jacobson et al., 2022). While traditionally thought to “coat” the Xi (Brockdorff et al., 1992; Brown et al., 1992; Clemson et al., 1996), 3D-SIM demonstrated Xist RNA to be distributed in foci (Cerase et al., 2014; Smeets et al., 2014; Sunwoo et al., 2015) with a preferential localization in regions of low DNA density (Smeets et al., 2014), matching the IC lacunae.

**Figure 4:**
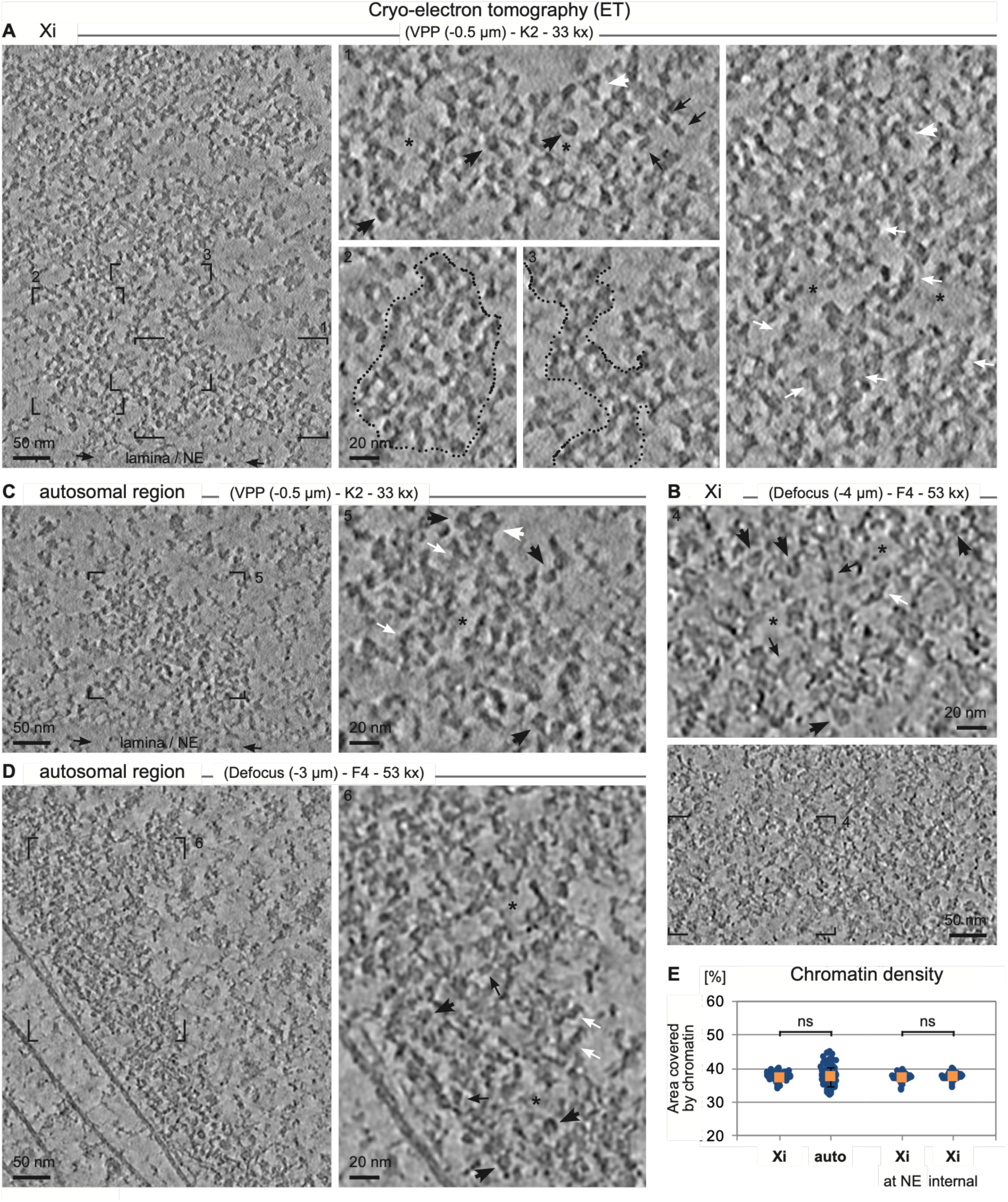
Heterochromatin organization is similar within the Xi and in autosomal regions. (A-D) Cryo-ET average slices (combined thickness of ∼5.5 nm; data collection parameters as indicated) of the Xi (A, B) and autosomal regions (C, D). Subareas showing heterochromatin are further magnified on the right / top. Asterisks indicate slightly enlarged interchromatin spaces. Black arrowheads and black arrows highlight disk views and side views of individual nucleosomes, respectively. White arrowheads and white arrows highlight zig-zag like conformations and short chains or stacks of nucleosomes, respectively. **(E)** Evaluation of heterochromatin density within the Xi and in autosomal regions (auto). For the Xi a comparison between heterochromatin at the nuclear envelope (NE) and internal regions is included. Mean = 37.3 nm / 37.4 nm / 37.2 nm / 37.4 nm. n = 88 / 118 / 39 / 49 measurements from 3 / 8 / 3 / 3 tomograms of 3 / 5 / 3 / 3 cells. Error bars represent standard deviation. Statistical significance based two-sided *t*-tests: p = 0.734 / p = 0.413.

Next, we compared the nuclear architecture of the Xi to autosomal chromatin regions (Fig 3, SFig 3D-E). In line with our APEX labeling data (Fig 1, SFig 2), the Xi was mainly heterochromatic (Fig 2, Fig 3A, SFig 3D), while autosomal regions were dominated by an extended IC and a mainly euchromatic / open chromatin organization (Fig 3B, SFig 3E), with only comparatively small heterochromatin domains (HCDs, arrowheads in Fig 3B). IC lacunae within the Xi were similar in size to HCDs in autosomal regions (Fig 3C). Because they usually extended beyond the field of view of the tomograms, we could not measure the length of the HCDs in the Xi and thus focused on their width alone. We found that HCDs in autosomal regions tended to be narrower compared to domains in the Xi and expectedly lacked the upper end of the range (Fig 3D). Next, we analyzed the dimensions of the IC. Within the Xi, IC lacunae exhibited a diameter of 90-570 nm when averaging length and width (Fig 3E, “size”).

Individual values for lacuna length (Fig 3E, “length & width”) reached up to 895 nm. Again, we could not obtain equivalent measurements of autosomal IC lacunae due to the expansive nature of the IC in autosomal regions. However, we evaluated the distance between the HCDs and found a similar range as the length and width measurements within the Xi (Fig 3E). We further found a significantly larger range for the width of the IC directly below the NPs in autosomal regions compared to the Xi, although the minimal values were similar (around 130 nm) (Fig 3F). Overall, these results confirm that compared to autosomal regions, the IC is reduced within the Xi and HCDs come closer together and form bigger clusters. At the same time, our measurements show that IC lacunae within the Xi can reach significant size. Further, the IC channels measured around 23 nm in width (Fig 3G) and thus allow the assumption that the passage of proteins from autosomal regions into the Xi and across the Xi is possible, in particular for smaller molecules like transcription factors but perhaps also for RNA polymerase II (Schultz et al., 1993; Bernecky et al., 2016). Taken together with the macromolecules we observed within the IC lacunae, these findings raise the question whether the active nuclear compartment (ANC) harboring functional nuclear processes (Cremer et al., 2020) within the Xi could be more similar to the ANC in autosomal regions than previously presumed from 3D SIM data (Smeets et al., 2014).

Finally, we examined how heterochromatin is associated with the NE. Heterochromatin is known to be anchored to the nuclear periphery, forming so called lamina-associated domains (LADs) (van Steensel and Belmont, 2017). Responsible for the anchoring are various proteins, including lamins and molecules that interact with chromatin directly or via binding proteins (Manzo et al., 2022; Shevelyov, 2023). The nuclear lamina, which lines the inner nuclear membrane, was visible in the cryo-electron tomograms in both en-face views (Fig 3H) as well as cross-sections (Fig 3A-B insets). In both the Xi and in autosomal regions, we regularly observed small protrusions of chromatin seemingly forming contacts with lamins and/or other proteins in the nuclear lamina (arrowheads in Fig 3A-B insets, Fig 3H), similar to what others described as well (Wang et al., 2025). Interestingly, the distance between heterochromatin and the NE was significantly smaller in autosomal regions than in the Xi (Fig 3I), potentially reflecting different mechanisms of anchoring heterochromatin to the nuclear lamina.

Altogether, our cryo-CLEM workflow allowed us to resolve the internal organisation of the Xi and its interaction with the NE in a native cellular context, adding on to existing work of the nuclear periphery (Mahamid et al., 2016; Wang et al., 2025) and providing a foundation for future studies of chromosome architecture and nuclear organization. In particular, the extent of functional processes taking place in the Xi and their exact spatial location requires further investigation.

### Cryo-ET provides insight into chromatin organization suggesting similar levels of compaction in heterochromatin domains inside and outside the Xi

Next, we analyzed the cryo-ET data to examine heterochromatin organization in more detail (Fig 4). Tomograms collected with the VPP (Fig 4A, C) resolved individual nucleosomes; both disk (black arrowheads) and side views (black arrows) were evident. Nucleosomes were also visible in data collected without the VPP (Fig 4B, D) but expectedly appeared less sharp due to the higher defocus applied. SFig 5 shows a side-by-side comparison of the different data collection modes, including a juxtaposition of cryo and APEX data. Nucleosomes regularly appeared to form short chains or stacks (Fig 4, white arrows) and occasionally seemed to adopt zig-zag-like conformations (white arrowheads) These assemblies seemingly contained only a few nucleosomes in the x/y plane but might extend further in 3D space. Within the HCDs nucleosomes were not distributed uniformly. Instead, we observed both regions with higher nucleosome density as well as areas with slightly enlarged interchromatin spaces (Fig 4A-D, asterisks). In most areas, chromatin was assembled into spherical patches of variable sizes (one example is outlined in Fig 4A inset 2), which presumably formed more complex networks in 3D. Occasionally chromatin adopted more elongated higher-order structures of variable shape and conformation (e.g. Fig 4A inset 3). Taken together, our data indicate that chromatin in RPE1 cells adopts irregular and variable arrangements without a recognizable preference for a specific higher order chromatin organization, which is in line with the majority of *in situ* chromatin studies (Eltsov et al., 2008; Ou et al., 2017; Cai et al., 2018; Hübner et al., 2022; Hou et al., 2023; Chen et al., 2025; Wang et al., 2025; Kreysing et al., 2026; Hou and Zhang, 2025; Kechagia and Medalia, 2026; Chen et al., 2026). Chromatin fibers were less distinct in our cryo data compared to our conventional APEX approach, presumably due to the APEX staining process (refer to (Ludwig, 2020)) enhancing the fiber structure. However, we did not observe regular fibers or fiber-like structures in the range of 30 nm with either of our approaches, neither within the Xi nor in autosomal regions, in contrast to previous *in vitro* studies of X chromosome chromatin organization (Naughton et al., 2010). Cell type specific differences could explain this contradiction, as fiber-like or elongated chromatin arrangements with diameters around 35 nm have been observed in certain cell types *in situ* (Hou et al., 2023; Kreysing et al., 2026), but not in others (Eltsov et al., 2008; Ou et al., 2017; Cai et al., 2018; Hübner et al., 2022; Chen et al., 2025; Wang et al., 2025).

Further, we found that chromatin density (percentage of area covered by nucleosomes) was not significantly different within the Xi compared to heterochromatin regions outside the Xi, nor in Xi domains close to the NE vs internal areas (Fig 4E). Together with the reduced IC in the Xi and previous findings that chromatin compaction levels of the Xi and its active counterpart are similar for small sub-chromosomal segments and only differ for larger ones (Naughton et al., 2010; Teller et al., 2011), this supports the hypothesis of the Cremer group (Teller et al., 2011) that the increased compaction of the Xi mainly stems from the reduced distance between the HCDs rather than from tighter packaging at the nucleosome level. Another study, however, reported differences in chromatin density based on ESI imaging on TEM sections (Baghestani, 2015).

Future studies will be needed to determine how chromatin organization and compaction in the Xi compare with those of autosomal facultative and constitutive heterochromatin, and to elucidate how mH2A-containing nucleosomes influence chromatin folding. Subtomogram averaging and template matching, which have recently been successfully applied to nucleosomes (Cai et al., 2018; Tan et al., 2023; Hou et al., 2023; Chen et al., 2025; Zhou et al., 2025; Wang et al., 2025; Kelley et al., 2026; Fatmaoui et al., 2026; Kreysing et al., 2026; Hou et al., 2026), will be instrumental in addressing these questions. Together with advances in *in situ* cryo-EM (Nogales and Mahamid, 2024; Majumder and Zhang, 2025; Subramaniam et al., 2026; Chen et al., 2026), these approaches are bringing structural biology into the cellular context, enabling molecular structures to be resolved while preserving their native environment.

## MATERIALS AND METHODS

### Conventional CLEM

For conventional CLEM (plastic embedding) with APEX2, sample preparation was performed based on (Martell et al., 2012; Lam et al., 2015) as described in (Ludwig, 2020). During the process, confocal fluorescence image stacks were acquired on a CorrSight confocal spinning disc microscope (FEI */* Thermo Fisher Scientific, Waltham, MA, USA) equipped with a cooling stage (FEI */* Thermo Fisher Scientific, Waltham, MA, USA and Julabo, Seelbach, Germany), Orca R2 CCD cameras (Hamamatsu Photonics, Hamamatsu City, Japan), a 40× oil objective (NA 1.3, EC Plan-Neofluar, Carl Zeiss, Oberkochen, Germany), 488 nm laser line, and standard filter sets. A H_2_O_2_ concentration of 0.5 mM and a reaction time of 5-10 min was used for the APEX2 reaction. Subsequently, transmission electron microscopy was performed on a Tecnai T12 (FEI / Thermo Fisher Scientific, Waltham, MA, USA) equipped with a Lab6 electron source and a 4k × 4k Eagle CCD camera (FEI / Thermo Fisher Scientific, Waltham, MA, USA) and operated at 120 kV. Sections were irradiated at low magnification for 10-20 min to prevent beam induced shrinkage during subsequent data acquisition and micrographs of the Xi were recorded at various magnifications. Finally, electron tomograms were collected on a Tecnai Arctica microscope (FEI / Thermo Fisher Scientific, Waltham, MA, USA) equipped with a high brightness field emission gun (S-FEG) and a Falcon 3EC 4k x 4k direct electron detector (Thermo Fisher Scientific, Waltham, MA, USA) and operated at 200 kV at room temperature. Dual-axis tilt series with manual rotation by 90° were recorded with Tomography Software (Thermo Fisher Scientific, Waltham, MA, USA) using a bidirectional tilt scheme starting at 0°, image acquisition at every second degree up to +/- 64 degrees, nanoprobe mode, objective aperture (100 µm), a C2 value of 40.5%, a dose of 20-25 e^-^Å^2^ per micrograph, a defocus of - 2 µm and a pixel size of 0.2787 nm (39k× nominal magnification). 15 tomograms from 10 different cells were collected at the Xi and 5 tomograms from 3 different cells outside the Xi at random autosomal regions.

### Cryo-CLEM

Overall, cryo-CLEM (sample preparation, cryo-FIB milling, data collection and data processing) was performed based on previously established protocols (Rigort et al., 2012; Schaffer et al., 2015; Zachs et al., 2020; Wagner et al., 2020; Tacke et al., 2021; Bieber et al., 2021). Details of our workflow are described below.

### Sample preparation for cryo-CLEM

Quantifoil or UltrAuFoil R2/2 Au200 grids (Quantifoil, Großlöbichau, Germany) were UV sterilized for at least 30 min in a biological safety cabinet (top side facing up). Then the grids were coated with Fibronectin (Sigma Aldrich / Merck, Darmstadt, Germany, #F1141) (1:400 in PBS (phosphate buffered saline), 1-4 h at 37°C, floating with top side down on drops on Parafilm), briefly washed by dipping them a few times into 5 successive drops of PBS (top side down), and transferred (with top side up) into a MatTek glass bottom dish (MatTek, Ashland, MA, USA; #P35G-1.5-14-C) also coated with Fibronectin and washed with PBS and containing 300 µl cell culture medium in the central glass area. As non-expressing cells tended to accumulate in the RPE1-eGFP-APEX2-mH2A cell lines over time, cells were subjected to fluorescence-activated cell sorting (FACS) on a BD FACSAria 3 sorter (BD Biosciences, Franklin Lakes, NJ, USA) in order to increase the number of cells suitable for CLEM on the grids. After sorting for the top 10-25% of eGFP-positive cells, the cultures were spun down and resuspended to a concentration of 80,000 cells / 150 µl. Taking care to prevent the grids from floating, 150 µl of medium were removed from the central glass area of the MatTek dish and replaced with 150 µl resuspended cell suspension. After ∼1.5 h medium was topped up to 2.5 ml total and the samples further incubated at least overnight. Nuclei were stained with Hoechst 33342 (Sigma Aldrich / Merck, Darmstadt, Germany, #B2261) (1 µg/ml, 1.5-2.5 h at 37°C) and the grids subsequently vitrified by plunge-freezing in liquid ethane on a Vitrobot Mark IV (Thermo Fisher Scientific, Waltham, MA, USA) using two filter papers (Electron Microscopy Sciences, Hatfield, PA, USA, #71166-65) on each side and the following settings: 37°C, 90-100% relative humidity, total blotting 1, wait and drain time 0 sec, blot force 8 or 9, blot time 3.5-9 sec depending on cell density. To improve targeting and correlation at later stages, 4 µl of fiducial marker beads (Life Technologies / Thermo Fisher Scientific, Waltham, MA, USA; #F8821, 1 µm Ø, red) (1:30 in PBS) were applied onto the cell-holding side of the grids just before plunge-freezing (Klumpe et al., 2021). Finally, the grids were clipped into CryoFIB autogrids (Thermo Fischer Scientific, Waltham, MA, USA, #1205101) of which the single dots as well as the double dot had been marked with two different colors of Sharpie permanent markers (Newell Brands, Atlanta, GA, USA) to facilitate grid orientation at later stages (Wagner et al., 2020).

### Cryo-fluorescence light microscopy

Light microscopic imaging under cryogenic conditions was performed on a CorrSight confocal spinning disk microscope (FEI / Thermo Fisher Scientific, Waltham, MA, USA) equipped with a cryo stage (FEI / Thermo Fisher Scientific, Waltham, MA, USA), Orca R2 CCD cameras (Hamamatsu Photonics, Hamamatsu City, Japan), 5x (NA 0.16) and 40x (NA 0.9) air objectives (both EC Plan-NEOFLUAR, Carl Zeiss, Oberkochen, Germany), 405 nm, 488 nm and 561 nm laser lines, and standard filter sets. For eGFP/488 nm imaging a GFP-specific filter was used. Image acquisition was performed in MAPS software (Thermo Fisher Scientific, Waltham, MA, USA). The 5x objective was used to collect overview images of the entire grid using transmission light to allow relocation of the cells on the cryo-FIB (see below). Image stacks of cells of interest were acquired with the 40x objective (163 nm pixel size). In widefield mode, stacks with 9 images and 0.5 µm z-distance were recorded for Hoechst, eGFP-mH2A, fiducial marker beads (red channel) and transmission light. In confocal spinning disk mode only eGFP-mH2A (488 nm) and fiducial marker beads (561 nm) were imaged using 61 images / stack and 0.2 µm z-distance. Only cells with intact and representative nuclei were pursued further.

### Target identification and cryo-FIB milling

The grids were transferred to an Aquilos DualBeam FIB-SEM (Thermo Fisher Scientific, Waltham, MA, USA) operated at 2 kV / 13 pA for the electron beam and 30 kV / 10 pA for the ion beam (for milling 30 pA to 1 nA was used; see below). After platinum coating (sputter coating (30 mA, 0.10-0.12 mbar, 15 sec) / GIS deposition (40-50 sec) / sputter coating again) target cells were relocated using MAPS 3.17 or 3.20 (Thermo Fisher Scientific, Waltham, MA, USA) by overlaying the newly acquired SEM images with the previously collected light microscopic data (maximum intensity projections). After cell identification at low magnification, SEM images of individual grid squares containing the target cells were acquired and used for precise correlation, making use of the fiducial marker beads as well as cellular features, in particular the nuclei. Target sites were selected and preparation for milling was performed in a semi-automatic way using AutoTEM 2.2 or 2.4 (Thermo Fisher Scientific, Waltham, MA, USA), manually choosing the milling angle (as low as possible, usually between 14 and 21 degrees stage tilt) as well as the final lamella position. Whenever possible we used 3D Correlation Toolbox (version 2.2.2) (Arnold et al., 2016; Klumpe et al., 2021; Bieber et al., 2021) at this step which allowed to calculate the z position of the marker beads in the fluorescence stack and thus improved targeting not only in xy but most importantly also in z direction. However, we consistently observed an offset between the target position identified by 3D Toolbox (as well as manual correlation) and the real position of the target (the Xi in our case) in 3D space. To compensate for this effect, we used the line tool in xT Microscope Control software (Thermo Fisher Scientific, Waltham, MA, USA) to measure 1.9 μm straight down from the suggested target location (black arrow in SFig 3A middle) and placed the final lamella position at the bottom end. Subsequently, step-wise milling (including stress relief cuts (Wolff et al., 2019) and using 4 milling steps) and initial thinning (polishing 1) was performed in AutoTEM with commonly used settings (Zachs et al., 2020; Wagner et al., 2020) down to a lamella thickness of ∼400 nm. The settings that worked best in our hands are listed in the supplementary methods. Further thinning (polishing 2) was performed manually in xT Microscope Control software with rectangular milling boxes using the 50 pA ion beam down to ∼300 nm thickness and the 30 pA ion beam to reach the final lamella thickness of ∼140-150 nm (confirmed by measuring with the line tool in xT Microscope Control software) as well as for a brief final polishing once all lamellae were completed. We did not use overtilt at any of the steps. Ultimately, samples were sputter coated with a thin layer of platinum (10 mA, 0.10-0.12 mbar, 3-4 sec).

### Cryo-electron tomography (ET) data collection

Cryo-ET data collection was performed on a Titan Krios G3 microscope (Thermo Fisher Scientific, Waltham, MA, USA) equipped with a high brightness field emission gun (X-FEG), 4k x 4k direct electron detector and post-column energy filter (K2 Summit / BioQuantum Gatan Imaging Filter (using a slit width of 15 eV) (both Gatan, Pleasanton, CA, USA) or Falcon 4i (F4i) / Selectris-X (using a slit width of 10 eV) (both Thermo Fisher Scientific, Waltham, MA, USA)), and operated at 300 kV in low dose mode. When loading the grids, the single dots of the CryoFIB autogrids were oriented up/down to ensure correct lamella orientation. Target areas were identified based on low to medium magnification micrographs guided by correlated overlay images of low magnification lamella overview images acquired during the final stages of FIB-milling with the corresponding fluorescence widefield images (maximum intensity projections). Tilt series were acquired with Tomography 5 Software (Thermo Fisher Scientific, Waltham, MA, USA) using a dose-symmetric tilt scheme (Hagen et al., 2017), pre-tilt of +/- 9 degrees, image acquisition at every third degree up to +/- 60 degrees from the start angle, a total dose per tomogram of 120-150 e^-^/Å^2^ collected as 6-8 dose fractions, and the following defocus values and pixel sizes: -5 or -6 µm defocus / 0.43 nm pixel size (K2 camera, 33k× nominal magnification), -4 or -6 µm / 0.47 nm (F4i, 26k×), -3, -4 or -6 µm / 0.24 nm (F4i, 53k×). With the K2 data was partially collected with Volta phase plate (VPP) (Danev et al., 2014; Fukuda et al., 2015), allowing us to reduce the total dose per tomogram to 100 e^-^/Å^2^ and defocus to -0.5 µm. With the F4i an objective aperture (70 µm) was used. Overall, we collected 6 tomograms at the Xi, 5 tomograms partially containing the Xi, and 21 tomograms (from 14 different cells) at random autosomal regions.

### Electron tomography (ET) data processing – cryo and conventional (APEX2)

For cryo-ET data, dose fractions were aligned with MotionCor2 (Zheng et al., 2017) and the resulting files sorted into order based on their tilt angles with a script generated in house. Using either the resulting combined .st file (for cryo-ET data) or directly inputting the tilt series .mrc files (for conventional ET data), all tomograms were reconstructed in IMOD (Kremer et al., 1996; Mastronarde and Held, 2017) using weighted back-projection and binning by 2 during the process. For conventional ET data, alignment was performed by tracking the gold fiducials and the two tilt axes were combined after tomogram generation. For cryo-ET data, patch tracking was used for alignment and the tomograms were denoised with CryoCare (Buchholz et al., 2019a; Buchholz et al., 2019b) after reconstruction. In Fiji/ImageJ (Schneider et al., 2012; Schindelin et al., 2015) all tomograms apart from those collected with Volta phase plate were then filtered to 2 nm using a Gaussian 3D filter and conventional ET data sets were binned again by 2 in all three directions. Final pixel sizes were 0.86 nm (K2) / 0.95 nm (F4i, 26k×) / 0.49 nm (F4i, 53k×) for cryo-ET data and 1.12 nm for conventional ET data. Groups of six (K2 and F4i, 26k×), eleven (F4i, 53k×) or five (conventional) consecutive tomographic slices were averaged to improve contrast, resulting in average slices of a combined thickness of ∼5.5 nm.

### Image segmentation and visualization

Image segmentation and 3D rendering was performed in Amira 3D version 2024.2 (Thermo Fisher Scientific, Waltham, MA, USA). AI Assisted Segmentation in the Segmentation+ Workroom was used to iteratively train and predict models (with VGG16, apart from ribosomes/macromolecules where shallow was used) for the various subcellular structures one by one. For nucleosomes and IC lacuna content several rounds of training were run with 90-100 epochs each and the labels and/or ROIs fixed and updated manually between the rounds as needed. For membranes 61 epochs were run and the result manually finalized. For ribosomes/macromolecules 20 epochs were used. To generate initial labels and to fix predicted output, the brush tool was used on denoised 8 bit data with appropriate thresholds applied (data was filtered to 1 pixel using a Gaussian 3D filter in Amira 3D for labeling membranes and ribosomes/macromolecules). For training and prediction, denoised 16 bit data, or for ribosomes/macromolecules the 16 bit rec file before denoising, were used. Small spots below an appropriate size (6 pixels in XY for nucleosomes and IC lacuna content, 4000 pixels in 3D for ribosomes/macromolecules, no filtering needed for membranes) were filtered out from the final segmentation result to reduce noise prior to generating the surface and applying the surface view.

### Image processing

Adobe Photoshop (Adobe Systems, San Jose, CA, USA) was used for stitching of overview images as well as post-acquisition image correlation and the generation of overlay images. Typically, distortion and sometimes minor warping of the fluorescence images was necessary for proper superimposition with the electron microscopic images. All other image processing was done in Fiji/ImageJ (Schneider et al., 2012; Schindelin et al., 2015), unless otherwise mentioned. Figures were prepared using Adobe InDesign (Adobe Systems, San Jose, CA, USA).

## ACKNOWLEDGEMENTS

Cryo-electron microscopy work and all data collection was performed at the NTU Institute of Structural Biology (NISB) Cryo-EM lab at Nanyang Technological University (NTU), Singapore. We are grateful to everyone at NISB, in particular Daniela Rhodes, Saw Wuan Geok and Ann Tran Bich Ngoc for their continuous support, Julien Lescar, Chong Wai Liew and Andrew Wong. We thank Zhang Li-Feng, Bilal Ahsan and Alexander Rigort for discussions and support. We are grateful to the Flow Cytometry Facility of the School of Biological Sciences (SBS) at NTU under Abdul Rashid BM Muzaki for the help with fluorescence activated cell sorting (FACS) sorting and to the Facility for Analysis, Characterisation, Testing and Simulation (FACTS) at NTU for support and housing the Titan Krios and Arctica microscopes. We thank Pei Yin Tan for performing experiments in the early stages of the project.

## AUTHOR CONTRIBUTIONS

BH and MLW performed all experiments. MLW performed conventional TEM imaging. HYSC performed data collection and reconstruction of low mag APEX2 tomograms. BH performed high mag APEX2- and all cryo-electron tomography data collection as well as all data analysis. AL, and in the early stages of the project also SS, conceived and supervised the work. BH and AL wrote the manuscript.

