## Supplementary Figures and Methods for "Native in situ architecture of the human inactive X chromosome revealed by correlative light and electron microscopy"

### Supplementary Information

### Supplementary Movies

**MovieS1:**

Segmentation and volume rendering of the Barr Body using AMIRA. Chromatin is shown in blue, IC lacuna are illustrated in magenta, macromolecular assemblies inside and outside the IC are shown in orange, ribosomes are in yellow, membranes are shown in white. Related to Figure 2.

### Supplementary Methods

#### Cloning

To generate the eGFP-APEX2-mH2A plasmid, human mH2A1 (macroH2A.1 histone; NCBI GenBank gene ID 9555) (kindly provided by Li-Feng Zhang) was inserted C-terminal of APEX2 into a Clontech eGFP-APEX2-C1 vector generated previously (Tan et al., 2020).

#### Cell culture

RPE1 cells (immortalized human retinal pigment epithelial cells) (kindly provided by Lu Lei) were cultured in DMEM/F12 medium (Gibco / Thermo Fisher Scientific, Waltham, MA, USA) supplemented with 10% FBS (fetal bovine serum) (CapricornScientific, Ebsdorfergrund, Germany) and penicillin-streptomycin (Gibco / Thermo Fisher Scientific, Waltham, MA, USA) and kept in humidified conditions at 37°C with 5% $CO_2$. Clonal cell lines expressing eGFP-APEX2-mH2A were generated according to standard protocols by transfecting the plasmid into RPE1 cells using Lipofectamine 3000 (Invitrogen / Thermo Fisher Scientific, Waltham, MA, USA) and adding geneticin/G418 (Gibco / Thermo Fisher Scientific, Waltham, MA, USA) (initially 500 mg/ml, later 200 mg/ml) to the cell culture medium during the selection period. These RPE1-eGFP-APEX2-mH2A cell lines were used for all experiments.

#### Data analysis and quantifications

Unless otherwise mentioned, measurements were performed in Fiji/ImageJ (Schneider et al., 2012; Schindelin et al., 2015) from tomogram average slices (compare ET data processing). Violin plots were generated in R (R Core Team; The RProject for Statistical Computing; R Foundation for Statistical Computing, Vienna, Austria; http://www.r-project.org/) using ggplot2 (Wickham, 2016) part of tidyverse (Wickham et al., 2019). All other graphs were generated in

Excel (Microsoft, Redmond, WA, USA) using Daniel's XL Tool box add-in (D. Kraus; www.xltoolbox.net) (version 7.2.) for bee swarm blots.

The thickness of chromatin fibers in the Xi was measured from conventional APEX2 ET data. An auto threshold (default method) was set for chromatin on one of the central average slices of the tomogram, "limit to threshold" selected in the measurements set up and the "area" measurement of the line tool (line width 1) used to determine fiber thickness. Only every third average slice was measured to avoid duplicates.

Dimensions of the interchromatin compartment (IC) and heterochromatin (HC) domains were measured from cryo-ET data. For measurements of IC channel width, only data collected with Volta phase plate (K2) or higher magnification (F4i, 53k×) was used. IC lacuna length represents the longest axis of the IC lacuna. For IC lacuna width, up to 5 measurements were taken per lacuna and subsequently averaged. IC lacuna size represents the average of length and averaged width. For the distance of HC domains in autosomal regions only the shortest distance between two domains was considered; thus this measurement represents minimal values. IC width at nuclear pores was measured between HC domains parallel to the nuclear envelope (NE). For tomograms in which the NE was close to perpendicular to the xy plane, measurements were performed in the center (largest diameter) of the nuclear pore. For tomograms in which the NE was cut at an angle, the IC lacuna below the nuclear pore was measured at its largest width.

The distance of chromatin from the nuclear envelope (NE) was measured perpendicular to the NE. To ensure this, tomograms were resliced to generate cross-section views of the NE (i.e. if the NE was running vertically, xz views were generated; if horizontally, yz views). If the NE was not oriented vertically in the resliced images, the resliced stack was rotated accordingly. Typically, different degrees of rotation had to be used for different areas within one tomogram. Rotated tomograms were resliced again to generate corrected xy views and average slices with a combined thickness of ~5.5 nm were generated as described under ET data processing. An auto threshold (default method) to highlight the IC was set based on a representative area including chromatin and the NE. "Limit to threshold" was selected in the measurements set up and the "area" measurement of the line tool (line width 1) used to determine the distance of chromatin from the NE. If pixels along the measurement line not representing the NE or chromatin (e.g. lamins) were spared out from the threshold, the brush tool was used to fill them prior to measuring.

Chromatin density was only evaluated from data collected with Volta phase plate (VPP) (K2) or higher magnification (F4i, 53k×). To ensure contrast settings to be as consistent across tomograms as possible, individual average slices (16 bit data) were segmented into seven chromatin density classes in R using the packages nucim (Schmid et al., 2017), bioimagetools (Schmid, V; http://volkerschmid.de, http://bioimaginggroup.github.io/bioimagetools/) and

EBImage (Pau et al., 2010) and a previously described algorithm (Markaki et al., 2012; Schmid et al., 2017). An all black image with the same dimensions as the input images was used as mask, x/y and z values were specified as 1:6 ratio and the beta factor was set to zero. Segmented images were combined back into stacks and particles smaller than 10 pixels filtered out using the “Analyze Particle” function in Fiji/ImageJ. Heterochromatin domains were then precisely outlined along the chromatin densities in the original average stack (excluding interchromatin compartment (IC) lacunas and channels as well as larger interchromatin spaces), the outline transferred to the corresponding binary segmented image and subsequently expanded by roughly half of the typical interchromatin space (3 pixels for VPP, 5 pixels for 53k×) to compensate for variable domain size and avoid bias due to the precise/tight outlining. A threshold was set to include the three densest classes comprising chromatin and overall area as well as the “area fraction” covered by chromatin measured using “limit to threshold”. Only every second average slice was measured to avoid duplicates.

**Hoechst staining**

RPE1-eGFP-APEX2-mH2A cells were seeded onto coverslips (no 1.5 thickness) coated with fibronectin (Sigma Aldrich / Merck, Darmstadt, Germany, #F1141) (1:400 in PBS (phosphate buffered saline), 1-4 h at 37°C) and allowed to settle at least overnight. Then nuclei were stained with Hoechst 33342 (Sigma Aldrich / Merck, Darmstadt, Germany, #B2261) (0.33 µg/ml, 2 h at 37°C) and the samples fixed with 4% PFA in PBS and mounted in 50% glycerol / 50% PBS. Confocal image stacks of the Hoechst and eGFP-mH2A signals were acquired on a CorrSight confocal spinning disk microscope (FEI / Thermo Fisher Scientific, Waltham, MA, USA) equipped with Orca R2 CCD cameras (Hamamatsu Photonics, Hamamatsu City, Japan), a 63x oil objective (NA 1.4, Plan-Apochromat, Carl Zeiss, Oberkochen, Germany), 405 nm and 488 nm laser lines, and standard filter sets. Pixel size was 103 nm and 200 nm z-distance was used. Corrections for chromatic shift were performed manually in Fiji/ImageJ (Schneider et al., 2012; Schindelin et al., 2015).

**Light microscopic analysis**

Unless otherwise mentioned, analyses were performed in Fiji/ImageJ (Schneider et al., 2012; Schindelin et al., 2015) from confocal image stacks of RPE1-eGFP-APEX2-mH2A cells stained with Hoechst. Graphs were generated as described in the main methods.
Localization of the Xi at the nuclear envelope (NE) was evaluated from fluorescent maximum intensity projections (MIPs) (2D) or from confocal image stacks (3D). Localization of the Xi at the nucleolus was determined from correlative data sets of eGFP-mH2A signals and transmission light images (2D) or from confocal image stacks (3D), identifying nucleoli based on experience from the transmission light or Hoechst images, respectively. For both

evaluations cells were counted as yes if either the DNA dense core of the Xi or the mH2A signals had contact with the NE / nucleolus.

**Measurement of macromolecule size**

Measurements were performed in Fiji/ImageJ from single tomogram slices of cryo-ET data. Only data collected with Volta phase plate (K2) or higher magnification (F4i, 53k×) was used. Graphs were generated as described in the main methods. For each molecule, measurements were taken on the slice with its largest dimensions or sharpest appearance. Molecule size was calculated as an average of its longest and shortest axis.

**Conventional electron tomography at lower magnification**

Electron tomograms of conventional CLEM samples at lower magnification (16k× nominal magnification) were collected as described in the main methods, however using microprobe mode, no objective aperture, a C2 value of 46.6%, a dose of 1.3 $e^{-}$Å$^{2}$ per micrograph, a defocus of -2 µm and a pixel size of 0.66 nm. 8 tomograms from 6 different cells were collected at the Xi. Processing was performed as described in the main methods apart from averaging only four consecutive tomographic slices (combined thickness of 10.48 nm; final pixel size was 2.62 nm).

**Electron tomogram reconstruction of cryo-ET data using SIRT**

Tomogram reconstruction using the simultaneous iterative reconstruction technique (SIRT) was performed as described in the main methods for using weighted back-projection, however SIRT with 8 iterations was selected on the Tomogram Generation page in IMOD (Kremer et al., 1996; Mastronarde and Held, 2017) and the tomogram after reconstruction filtered to only 1 nm using a Gaussian 3D filter in Fiji/ImageJ. SIRT tomograms were not denoised with CryoCare.

**Settings for cryo-FIB milling (AutoTEM)**

| | |
|---|---|
| Lamella dimensions: | 10 µm depth, typically 10 µm width (down to 8 µm if needed); 150 nm final thickness |
| Front/rear windows: | 4.5 µm height |
| Correction factor: | 0.5 |
| Stress relief cuts: | 10 µm height, 0.75 µm width, typically 1 nA ion beam current (0.3-0.5 nA for very thin cells/areas), 400% depth correction |
| Milling: | rough: 1 nA ion beam current, 1.3 µm offset (resulting in 2.75 µm lamella thickness), 60% depth correction |
| | medium: 0.5 nA, 650 nm offset (1.45 µm), 200% |

| | |
|---|---|
| | fine: 0.3 nA, 350 nm offset (850 nm), 200% |
| | finer: 0.1 nA, 200 nm offset (550 nm), 200%, 60 sec DCM interval |
| Thinning: | polishing 1: 50 pA, 125 nm offset (400 nm), 200% |
| | (polishing 2: done manually, refer to main methods) |
| Overtilt: | 0° for all steps |

HFW (horizontal field width): 140 µm for all electron beam images

For all other settings default values were used.

### Supplementary Figures

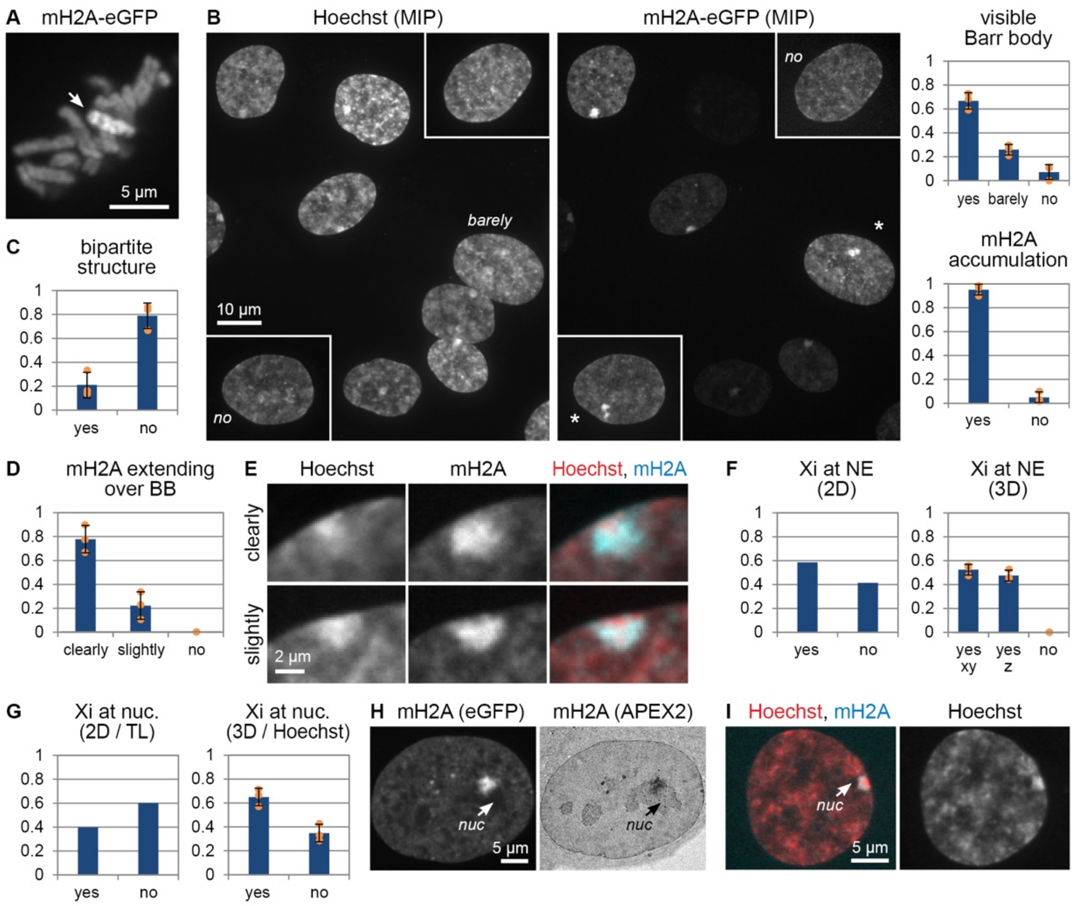


**SFigure 1: Light microscopic analysis of the mH2A-eGFP-APEX2 RPE1 cell line. (A)** Metaphase cell (mH2A-eGFP signals) with the Xi highlighted by the arrow (maximum intensity projection). **(B)** Maximum intensity projections of nuclei (DNA) stained with Hoechst (left) and mH2A-eGFP signals (middle). Right: Evaluation of the frequency of cells exhibiting a visible Barr body (i.e. DNA dense core of the Xi) based on Hoechst signals (top) or sites of mH2A accumulation (bottom). Mean = 0.67 / 0.26 / 0.07 and 0.95 / 0.05, respectively. n = 68 and 162 cells, respectively, from 3 subpopulations. Error bars represent standard deviation. **(C)** Evaluation of the frequency of cells showing a bipartite structure of the Xi based on mH2A-eGFP signals (as in the cells marked by asterisks in (B)). Mean = 0.21 / 0.79. n = 408 cells (68 evaluated in 3D, 340 in 2D) from 3 subpopulations (3 in 3D, 1 in 2D). Error bars represent standard deviation. **(D)** Evaluation of the frequency of cells exhibiting mH2A-eGFP signals extending over the Barr body (BB). Mean = 0.78 / 0.22 / 0. n = 68 cells from 3 subpopulations. Error bars represent standard deviation. **(E)** Representative examples of mH2A-eGFP signals extending over the BB as evaluated in (D). **(F)** Evaluation of the frequency of cells in which the Xi is located at the nuclear envelope (NE) evaluated either in 2D (left) or in 3D (right). Yes xy = Xi at NE in a single light optical section. Yes z = Xi at NE after examination of the confocal image stack. Mean = 0.59 / 0.41 and 0.52 / 0.48 / 0,

respectively. n = 340 and 68 cells, respectively, from 1 and 3 subpopulations. Error bars represent standard deviation. **(G-I)** Evaluation of the frequency of cells in which the Xi is located at a nucleolus (nuc) (G), evaluated either in 2D (left) based on transmission light (TL) images as shown in (H; confocal light optical section / TEM image) or in 3D (right) based on Hoechst staining as shown in (I; confocal light optical section). Mean = 0.40 / 0.60 and 0.65 / 0.35, respectively. n = 289 and 53 cells, respectively, from 1 and 3 subpopulations. Error bars represent standard deviation.

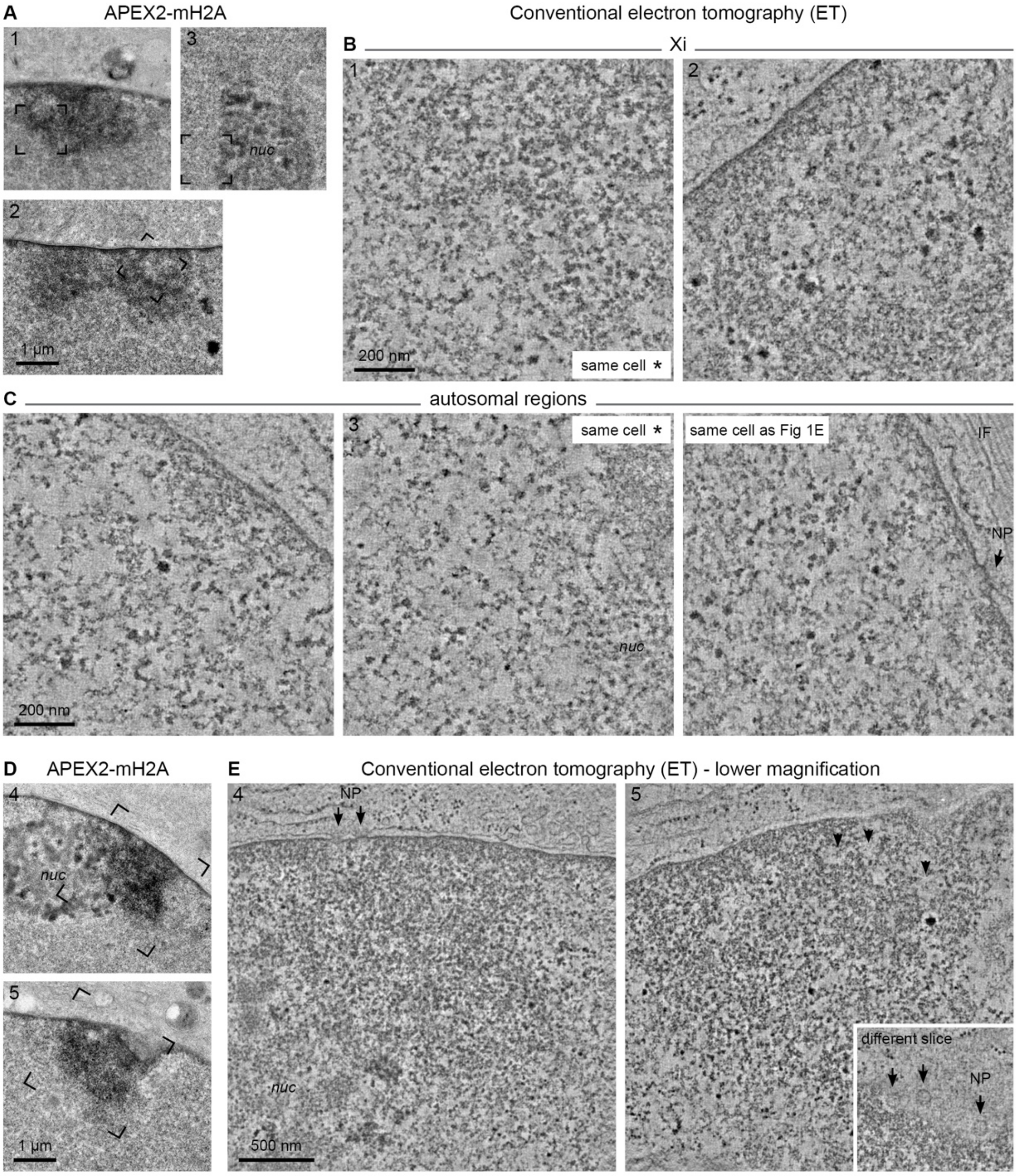


**SFigure 2: Conventional CLEM and ET of the Xi and autosomal regions. (A, D)** TEM images of APEX2-mH2A signals (1-2, 4-5) or an autosomal region including a nucleolus (nuc) (3). **(B, C)** ET average slices (combined thickness of 5.6 nm; -2 µm defocus, 39k×) of the boxed areas in (A) and two additional examples for autosomal regions. Top row: Xi. Bottom row: autosomal regions. NP = nuclear pore. IF = intermediary filaments. Tomograms collected from the same cells are indicated. **(E)** ET average slices (combined thickness of 10.5 nm; -2 µm defocus, 16k×) of the boxed areas in (D). Arrowheads point at IC lacunae beneath nuclear pores (NP) (see inset).

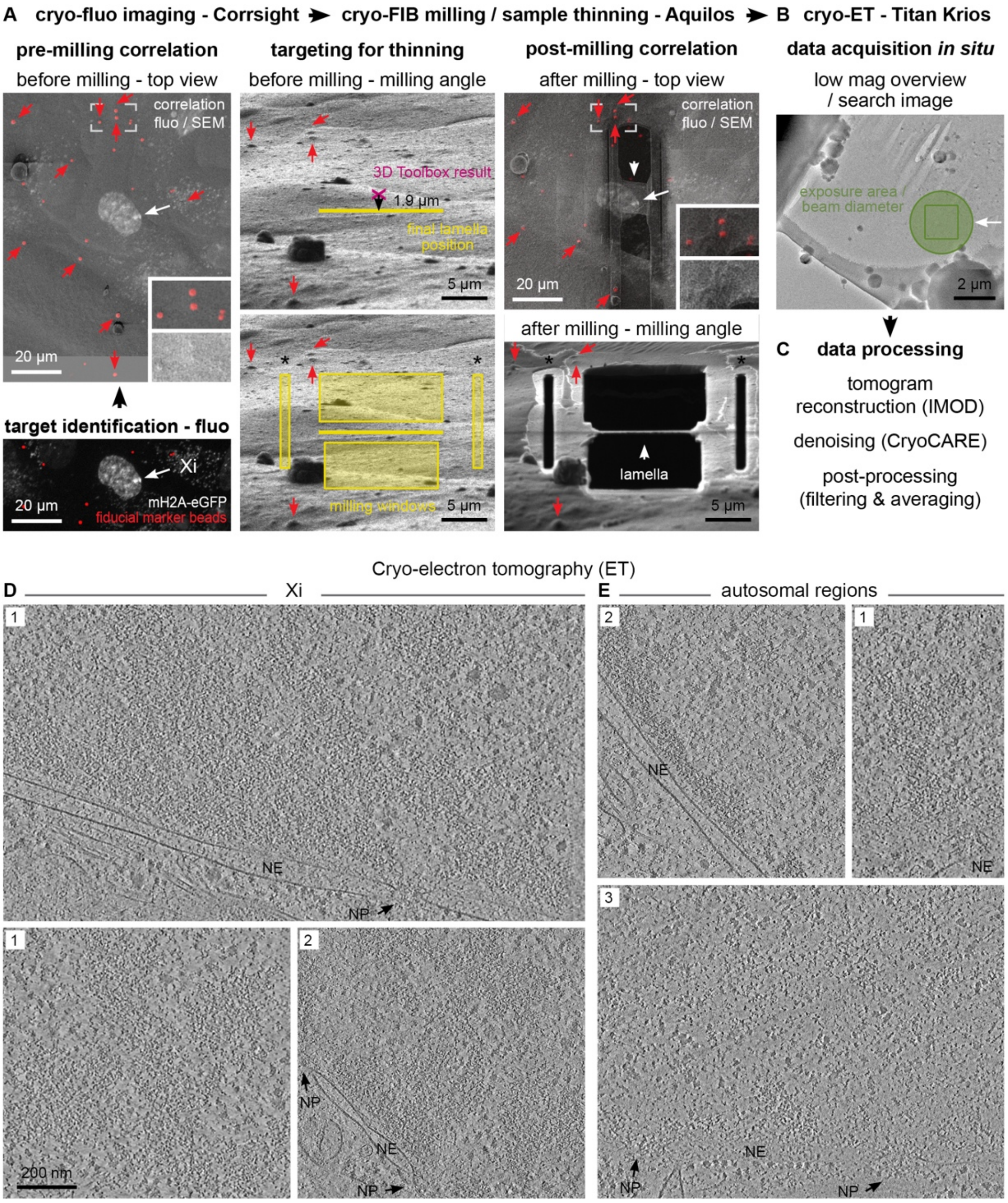


**SFigure 3: Cryo-CLEM workflow and cryo-ET of the Xi and autosomal regions. (A-C)** Cryo-CLEM workflow. Cells were grown on grids and fiducial marker beads applied just before plunge-freezing in liquid ethane on a Vitrobot (not shown). (A) Two-step LM and EM correlation for Xi targeting making use of the marker beads. Cryo-fluorescence images of target cells (arrow pointing at the Xi) were collected on the Corrsight (left, bottom; confocal spinning disk mode, maximum intensity projection; gray = mH2A-eGFP signals, red = fiducial marker beads) and overlaid with the SEM images from the Aquilos cryo-FIB for pre-milling correlation for correct xy targeting (left, top). Red arrows indicate the position of marker beads. The boxed area is magnified in the inset and includes also the SEM image alone (bottom inset), showing the bumps of the marker beads. At the milling angle (middle), the final lamella position was defined by using 3D Correlation Toolbox whenever possible to

improve targeting in z, measuring 1.9 µm straight down (black arrow) from the suggested target location (magenta), and placed the final lamella position at the bottom end (yellow). Milling windows and stress relief cuts (*) (light yellow) were placed accordingly. The milled result is shown on the bottom right. Post-milling correlation (right, top) was performed equivalent to pre-milling correlation using low mag SEM images of the milled lamella (arrowhead). (B) Based on the post-milling correlation the region of interest for cryo-ET data collection was determined on the Titan Krios. The green circle indicates the beam diameter, the square the exposure area. (C) Main steps of data processing. Refer to methods for details. **(D-E)** Cryo-ET average slices (combined thickness of ~5.5 nm) of the Xi (D) and autosomal regions (E). Numbers indicate data collection parameters: 1 = F4i, -6 µm defocus, 26k×; 2 = F4i, -3 µm defocus, 53k×; 3 = VPP, K2, -0.5 µm defocus, 33k×). NE = nuclear envelope. NP = nuclear pore.

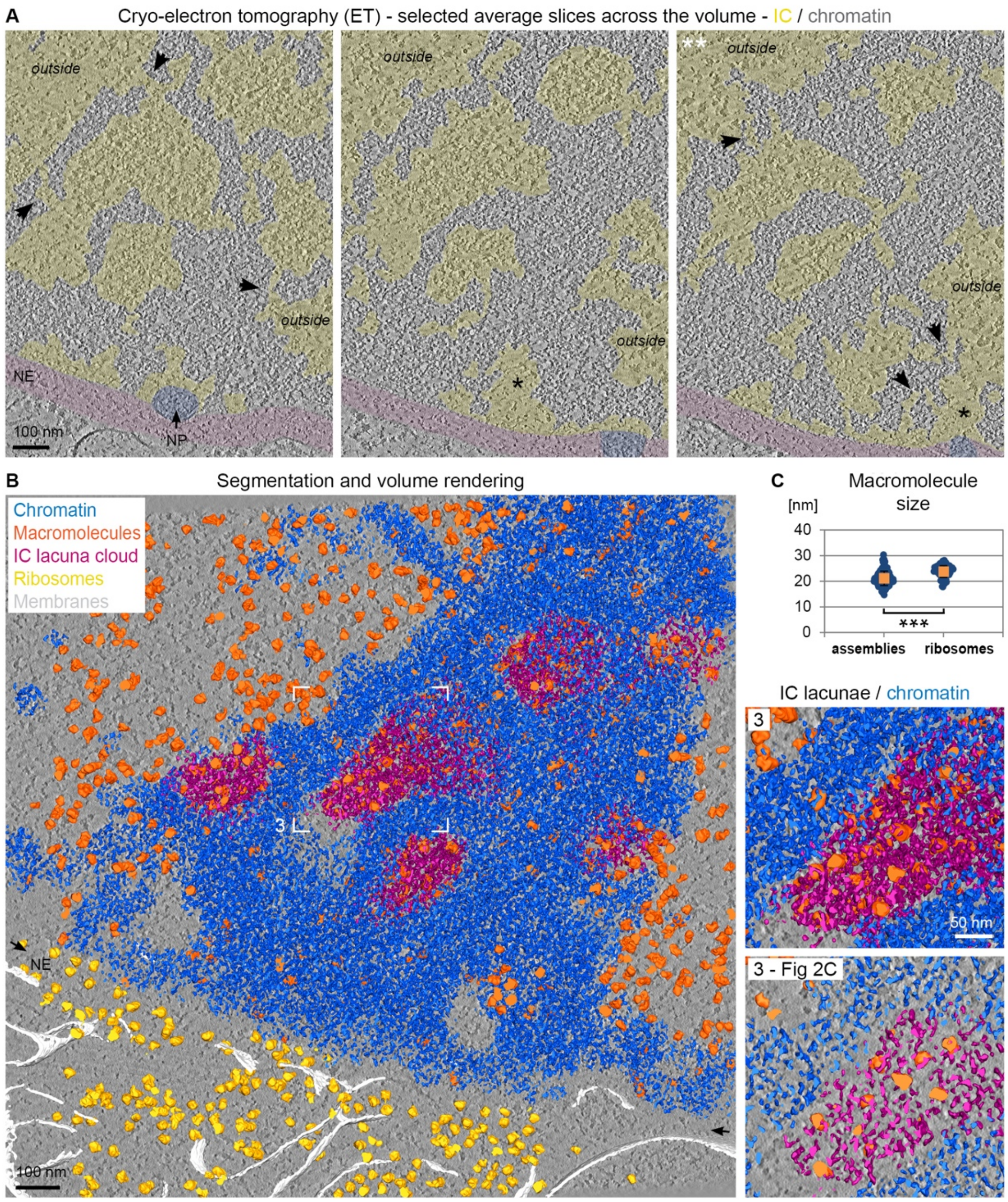


**SFigure 4: The IC network in the Xi as revealed by cryo-ET and full volume segmentation and rendering. (A)** Selected cryo-ET average slices across the volume of the tomogram shown in Fig 2B (combined thickness of 5.2 nm each; VPP, K2, -0.5 µm defocus, 33k×). The right image marked with ** is the same as shown in Fig 2B. IC/EC overlaid with yellow. NE = nuclear envelope, overlaid with magenta. NP = nuclear pore, overlaid with purple. Arrowheads point at IC channels connecting the IC lacunae within the Xi with the nucleoplasm outside the Xi. Asterisks highlight IC lacunae below nuclear pores. **(B)** Segmentation and volume rendering corresponding to Fig 2C but comprising a thickness of 112.3 nm. A single tomogram slice is shown in the background (gray). The boxed area is further magnified on the right. The bottom inset shows the corresponding small volume rendering (20.7 nm) of boxed area 3 in Fig 2C. NE = nuclear envelope (orientation indicated by black arrows). Blue = chromatin, orange = macromolecules, magenta = IC lacuna cloud,

yellow = ribosomes, white = membranes. Note that the NE could not be segmented due to lack of contrast as a consequence of being cut at an angle. **(C)** Evaluation of the size of macromolecular assemblies in the Xi IC lacunae (assemblies) and of ribosomes. Mean = 21.1 nm / 23.8 nm. n = 140 / 84 measurements from 3 tomograms of 3 cells. Error bars represent standard deviation.

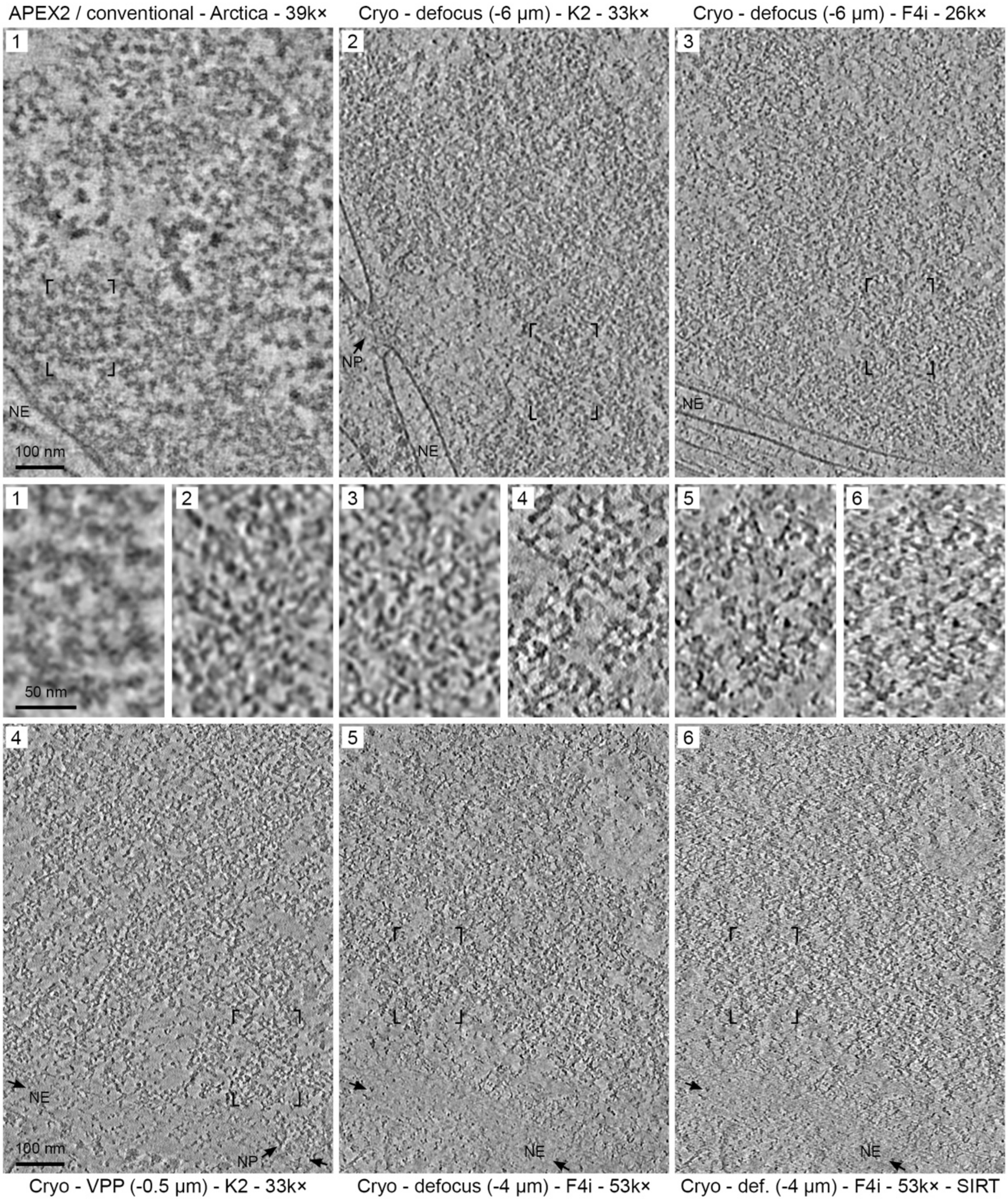


**SFigure 5: Comparison of ET data collection modes.** Data collection parameters are indicated. The boxed areas in the top and bottom row are magnified in the middle. Data set 1 is conventional sample preparation with APEX2, data set 2-6 are cryo samples. Data set 1-5 were reconstructed using weighted back-projection, data set 6 is the same as data set 5 but reconstructed using the simultaneous iterative reconstruction technique (SIRT). ET average slices with a combined thickness of ~5.5 nm are shown. NE = nuclear envelope (where applicable orientation indicated by black arrows). NP = nuclear pore.
